# Revealing neuro-computational mechanisms of reinforcement learning and decision-making with the hBayesDM package

**DOI:** 10.1101/064287

**Authors:** Woo-Young Ahn, Nathaniel Haines, Lei Zhang

**Affiliations:** Department of Psychology, The Ohio State University, Columbus, OH; Institute for Systems Neuroscience, University Medical Center Hamburg-Eppendorf, Hamburg, Germany

**Keywords:** Reinforcement learning, decision-making, hierarchical Bayesian modeling, model-based fMRI

## Abstract

Reinforcement learning and decision-making (RLDM) provide a quantitative framework and computational theories, with which we can disentangle psychiatric conditions into basic dimensions of neurocognitive functioning. RLDM offer a novel approach to assess and potentially diagnose psychiatric patients, and there is growing enthusiasm on RLDM and Computational Psychiatry among clinical researchers. Such a framework can also provide insights into the brain substrates of particular RLDM processes as exemplified by model-based functional magnetic resonance imaging (fMRI) or electroencephalogram (EEG). However, many researchers often find the approach too technical and have difficulty adopting it for their research. Thus, there remains a critical need to develop a user-friendly tool for the wide dissemination of computational psychiatric methods. We introduce an R package called hBayesDM (*<u>h</u>*ierarchical *<u>Bayes</u>*ian modeling of *<u>D</u>*ecision-*<u>M</u>*aking tasks), which offers computational modeling on an array of RLDM tasks and social exchange games. The hBayesDM package offers state-of-the-art hierarchical Bayesian modeling, where both individual and group parameters (i.e., posterior distributions) are estimated simultaneously in a mutually constraining fashion. At the same time, it is extremely user-friendly: users can perform computational modeling, output visualization, and Bayesian model comparisons–each with a single line of coding. Users can also extract trial-by-trial latent variables (e.g., prediction errors) required for model-based fMRI/EEG. With the hBayesDM package, we anticipate that anyone with minimal knowledge of programming can take advantage of cutting-edge computational modeling approaches and investigate the underlying processes of and interactions between multiple decision-making (e.g., goal-directed, habitual, and Pavlovian) systems. In this way, it is our expectation that the hBayesDM package will contribute to the dissemination of advanced modeling approaches and enable a wide range of researchers to easily perform computational psychiatric research within their populations.

## 1. Introduction

Computational modeling (a.k.a., cognitive modeling) describes human information processing with basic principles of cognition, which are defined in formal mathematical notations (Figure 1). Unlike verbalized or conceptualized approaches, computational modeling has the merit of allowing researchers to generate precise predictions and quantitatively test competing hypotheses (Busemeyer & Diederich, 2010; Forstmann & Wagenmakers, 2015; Lewandowsky & Farrell, 2010). Computational modeling has been particularly useful in reinforcement learning and decision-making (RLDM) fields (Dayan & Daw, 2008; Rangel, Camerer, & Montague, 2008); computational modeling has also been integrated into the analysis of neural data including functional magnetic resonance imaging (fMRI) and electroencephalogram (EEG) (e.g., Cavanagh, Eisenberg, Guitart-Masip, Huys, & Frank, 2013; Daw, O'Doherty, Dayan, Seymour, & Dolan, 2006; Gläscher, Hampton, & O'Doherty, 2009; Hampton, Bossaerts, & O'Doherty, 2006; Iglesias et al., 2013; Li, Schiller, Schoenbaum, Phelps, & Daw, 2011; Mars et al., 2008; O'Doherty, Hampton, & Kim, 2007; O'Doherty, Dayan, Schultz, & Deichmann, 2004; Xiang, Lohrenz, & Montague, 2013).

**Figure 1.**
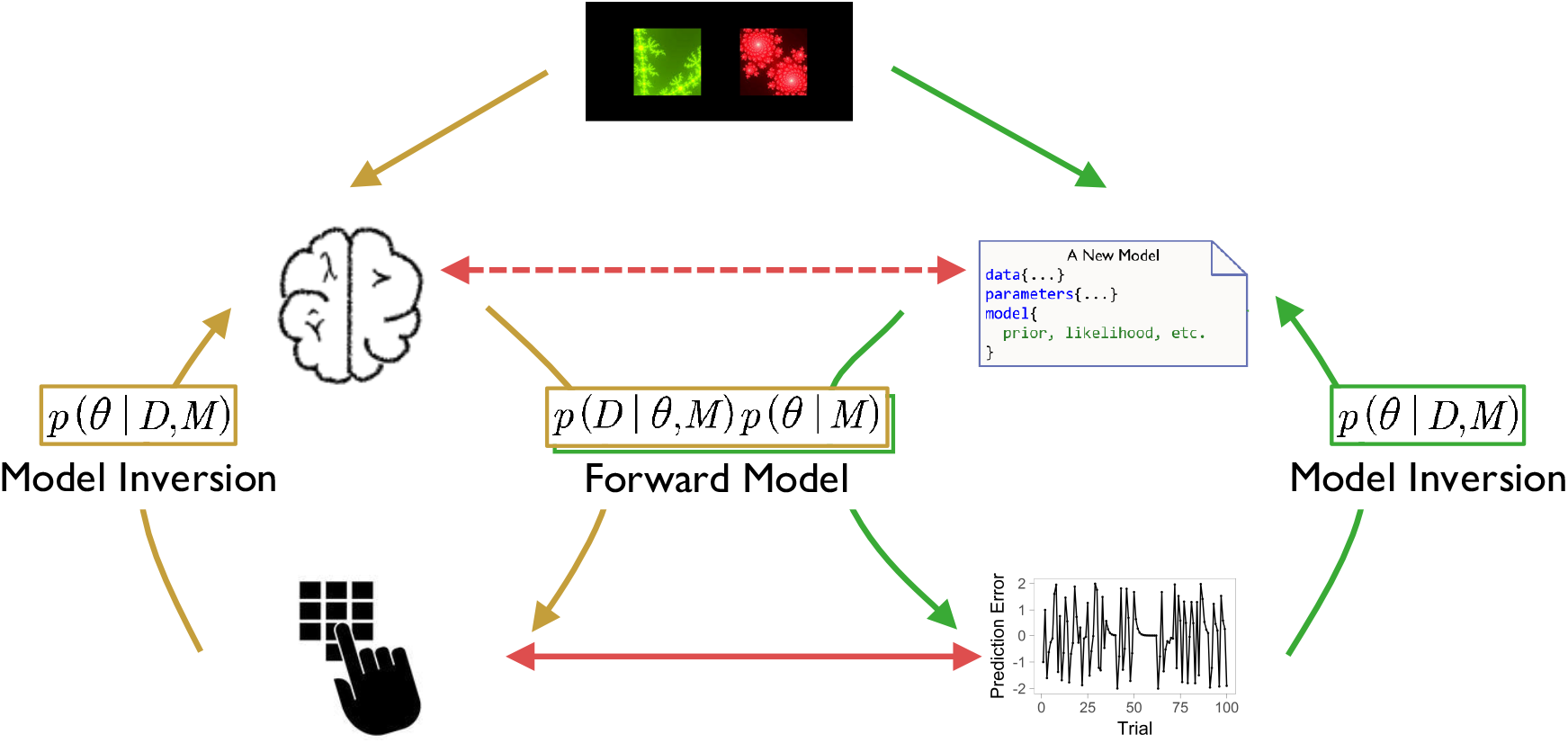
Conceptual schema of computational modeling. Starting with a certain RLDM paradigm, the left pathway (yellow arrows) represents that the human brain produces behavioral responses (forward model) that we observe and measure. These observed outcomes can be used to make inference on cognitive mechanisms (model inversion), but oftentimes this is difficult to achieve. One solution is to build cognitive models (green arrows) that produce model predictions (forward model) and can also be inferred by those predictions (model inversion). As such, we are able to approximate brain mechanisms (dashed red line) by directly linking model predictions (e.g., reward prediction error) with observed outcomes (solid red line).

As summarized in recent review papers (Ahn & Busemeyer, 2016; Friston, Stephan, Montague, & Dolan, 2014; Huys, Maia, & Frank, 2016; Montague, Dolan, Friston, & Dayan, 2012; Stephan, Bach, et al., 2016a; Stephan, Binder, et al., 2016b; Stephan, Iglesias, Heinzle, & Diaconescu, 2015; X.-J. Wang & Krystal, 2014; Wiecki, Poland, & Frank, 2015), computational modeling has gained much attention for its usefulness in investigating psychiatric conditions. Exemplified by the Research Domain Criteria (RDoC; Insel, 2014) and Precision Medicine, there is also a growing consensus that diagnosis and treatment decisions should incorporate underlying cognitive and neurobiological underpinnings of psychiatric conditions instead of relying only on behavioral symptoms. To this end, a new field called *Computational Psychiatry* (e.g., Friston et al., 2014; Montague et al., 2012) aims to discover neurocognitive mechanisms underlying normal and abnormal conditions by combining cutting-edge neurobiological and computational tools.

Performing computational psychiatric research, however, especially computational modeling, is a challenging task for many clinical researchers or those with limited quantitative skills. Computational modeling involves multiple steps including designing/adopting laboratory tasks, building a theoretical framework of the task with a set of assumptions and mathematical equations, formulating multiple computational models based on the assumptions, estimating model parameters of each model, and quantitatively comparing the models of interest (e.g., Busemeyer & Diederich, 2010; Wiecki et al., 2015). It is a pressing issue how to train clinical researchers in mental health (e.g., psychiatrists and clinical psychologists) so that they can receive in-depth training across several related fields including cognitive science, advanced statistics, and neuroscience (Montague et al., 2012). For the dissemination of Computational Psychiatry, we believe there remains a critical need to develop user-friendly tools for computational modeling. In fact, there exist several software packages but most of them are for a single class of modeling such as sequential sampling models (Matzke et al., 2013; Singmann et al., 2016; Vincent, 2015; Wabersich & Vandekerckhove, 2014; Wiecki, Sofer, & Frank, 2013). An exception is the Variational Bayesian Analysis (VBA) MATLAB toolbox (Daunizeau, Adam, & Rigoux, 2014), which allows users to fit and compare various models with variational Bayesian algorithms. However, we believe users still need some amount of programming skills and background in computational modeling to model various tasks with the VBA toolbox.

In this article, we describe a free R package, ***hBayesDM*** (*<u>h</u>*ierarchical *<u>Bayes</u>*ian modeling of *<u>D</u>*ecision-*<u>M</u>*aking tasks), which we developed for the dissemination of computational modeling to a wide range of researchers. The hBayesDM package offers hierarchical Bayesian analysis (HBA; see Section 3 for more details about HBA) of various computational models on an array of decision-making tasks (see **Table 1** for a list of tasks and models currently available). With the user-friendly hBayesDM package, users can perform model fitting with HBA, output visualization, and model comparisons – *each with a single line of coding*. Example datasets are also available to make it easy to use hBayesDM. Users can also extract trial-by-trial latent variables (e.g., prediction errors) that are required for model-based fMRI/EEG (see Section 5.5). Experienced users can also develop new models based on our framework and codes. All source codes are publicly available at our GitHub repository (https://github.com/ccs-lab/hBayesDM). Users can ask questions to our mailing list (https://groups.google.com/forum/#!forum/hbayesdm-users) or make suggestions by posting new issues to the GitHub repository. By making all steps for computational modeling user-friendly, we hope the hBayesDM package will allow even researchers with minimal programming knowledge to perform certain computational psychiatric research.

The remainder of this article is organized as follows. First, we describe the list of tasks and models that are currently implemented in the hBayesDM package (Section 2). Second, we briefly describe HBA and why we adopt HBA for computational modeling (Section 3). Third, we explain the detailed mathematical formulation of hierarchical Bayesian models (Section 4). Fourth, we provide step-by-step tutorials on how to use the hBayesDM package (Section 5). Lastly, we discuss future directions and potential limitations of the package (Section 6). Readers who are not interested in the technical details may skip Section 3 and equations in Section 4.

## 2. Tasks and computational models implemented in hBayesDM

**Table 1** shows the list of tasks and computational models currently implemented in the hBayesDM package (as of version 0.2.3.2). Note that some tasks have multiple computational models and users can compare the model performance within the hBayesDM framework (see Section 5). To fit models to a task, prepare trial-by-trial data as a text file (*.txt) where each row (observation) contains the columns required for the given task (see **Table 1**). Users can also use its sample dataset as an example.

Below, we describe each task and its computational model(s), briefly review its applications to healthy and clinical populations, and describe model parameters. For brevity, we refer readers to original papers for the full details of experimental design and computational models, and to the package help files for example codes that detail how to estimate and extract parameters from each model. Package help files can be found by issuing the following command within the R console:

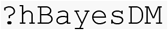

The above command will open the main help page, where one can then navigate to the corresponding task/model. Users can also directly look up a help file for each task/model by calling its help file, which follows the form `?function_name` (e.g., `?dd_cs`, see **Table 1** for a list of functions). Each help file provides working codes to run a concrete real-data example from start to finish.

### 2.1. The Delay Discounting task

The Delay Discounting task (DDT; Rachlin, Raineri, & Cross, 1991) is designed to estimate how much an individual discounts temporally delayed larger outcomes in comparison to smaller-sooner ones. On each trial of the DDT, two options are presented: a sooner and smaller reward (e.g. $5 now) and a later and larger reward (e.g. $20 next week). Subjects are asked to choose which option they prefer on each trial.

The DDT has been widely studied in healthy populations (e.g., Green & Myerson, 2004; Kable & Glimcher, 2007) and DD has been associated with cognitive abilities such as intelligence (Shamosh et al., 2008) and working memory (Hinson, Jameson, & Whitney, 2003). Steeper delay discounting is a strong behavioral marker for addictive behaviors (Ahn & Vassileva, 2016; Ahn, Ramesh, Moeller, & Vassileva, 2016; Bickel, 2015; Green & Myerson, 2004; MacKillop, 2013) and has also been associated with other psychiatric conditions including schizophrenia (Ahn, Rass, Fridberg, Bishara, Forsyth, Breier, et al., 2011b; Heerey, Matveeva, & Gold, 2011; Heerey, Robinson, McMahon, & Gold, 2007) and bipolar disorder (Ahn, Rass, Fridberg, Bishara, Forsyth, Breier, et al., 2011b). The hBayesDM package currently contains three different models for the DDT:

1. `dd_cs` (Constant Sensitivity model, Ebert & Prelec, 2007)
  Exponential discounting rate ( 0 < *r* <1)
  Impatience ( 0 < *s* <10)
  Inverse temperature ( 0 < *β* < 5)
2. `dd_exp` (Exponential model, Samuelson, 1937)
  Exponential discounting rate ( 0 < *r* <1)
  Inverse temperature ( 0 < *β* < 5)
3. `dd_hyperbolic` (Hyperbolic model, Mazure, 1987)
  Discounting rate ( 0 < *k* <1)
  Inverse temperature ( 0 < *β* < 5)

#### 2.1.1 DDT: Parameter Descriptions

In exponential and hyperbolic models, temporal discounting of future (delayed) rewards is described by a single parameter, discounting rate ( 0 < *r* < 1), which indicates how much future rewards are discounted. High and low discounting rates reflect greater and lesser discounting of future rewards, respectively. In exponential and hyperbolic models, the value of a delayed reward is discounted in an exponential and hyperbolic form, respectively. The Constant-Sensitivity (CS) model has an additional parameter called time-sensitivity (0 < *s* < 10). When *s* is equal to 1, the CS model reduces to the exponential model. Values of *s* near 0 lead to a simple “present-future dichotomy” where all future rewards are steeply discounted to a certain subjective value, irrespective to delays. Values of *s* greater than 1 result in an “extended-present” heuristic, in which rewards during the extended present are valued nearly equally and future rewards outside the extended present have zero value.

All models use the softmax choice rule with an inverse temperature parameter (Kaelbling, L. P., Littman, & Moore, 1996; Luce, 1959), which reflects how deterministically individuals’ choices are made with respect to the strength (subjective value) of the alternative choices. High and low inverse temperatures represent more deterministic and random choices, respectively.

### 2.2. The Iowa Gambling task

The Iowa Gambling task (IGT, Bechara, Damasio, Damasio, & Anderson, 1994) was originally developed to assess decision-making deficits of patients with ventromedial prefrontal cortex lesions. On each trial, subjects are presented with four decks of cards. Two decks are advantageous (good) and the other two decks disadvantageous (bad) in terms of long-term gains. Subjects are instructed to choose decks that maximize long-term gains, which they are expected to learn by trial and error. From a statistical perspective, the IGT is a four-armed bandit problem.

The IGT has been used extensively to study decision-making in several psychiatric populations (Ahn et al., 2014; Bechara & Martin, 2004; Bechara et al., 2001; Bolla et al., 2003; Grant, Contoreggi, & London, 2000; Vassileva, Gonzalez, Bechara, & Martin, 2007). The hBayesDM package currently contains three different models for the IGT:

1. `igt_pvl_decay` (Ahn et al., 2014; Ahn, Krawitz, Kim, Busemeyer, & Brown, 2011a)
  Decay rate ( 0 < *A* <1)
  Shape ( 0 < *α* < 2)
  Consistency ( 0 < *c* < 5)
  Loss Aversion ( 0 < *λ* <10)
2. `igt_pvl_delta` (Ahn, Busemeyer, Wagenmakers, & Stout, 2008)
  Learning rate ( 0 < *A* <1)
  Shape ( 0 < *α* < 2)
  Consistency ( 0 < *c* < 5)
  Loss Aversion ( 0 < *λ* <10)
3. `igt_vpp` (Worthy, Pang, & Byrne, 2013)
  Learning rate ( 0 < *A* <1)
  Shape ( 0 < *α* < 2)
  Consistency ( 0 < *c* < 5)
  Loss Aversion ( 0 < *λ* <10)
  Perseverance gain impact (−∞<*ε*_*p*_ <∞)
  Perseverance loss impact ( −∞<*ε*_*n*_ <∞)
  Perseverance decay rate ( 0 < *k* <1)
  Reinforcement learning weight ( 0 < *ω* <1)

#### 2.2.1 IGT: Parameter Descriptions

The prospect-valence learning (PVL) model with delta rule (PVL-delta) uses a Rescorla-Wagner updating equation (Rescorla & Wagner, 1972) to update the expected value of the selected deck on each trial. The expected value is updated with a learning rate parameter (0 < *A* < 1) and a prediction error term, where *A* close to 1 places more weight on recent outcomes and *A* close to 0 places more weight on past outcomes; the prediction error is the difference between predicted and experienced outcomes. The shape ( 0 < *α* < 2) and loss aversion ( 0 < *λ* < 10) parameters control the shape of the utility (power) function and the effect of losses relative to gains, respectively. Values of *α* greater than 1 indicate that the utility of outcome is convex, and values less than 1 indicate that the utility is concave. Values of *λ* greater than and less than 1 indicate greater or reduced sensitivity to losses relative to gains, respectively. The consistency parameter ( 0 < *c* < 5) is an inverse temperature parameter (refer to section 2.1.1 for details).

The PVL model with decay rule (PVL-decay) contains the same shape, loss aversion, and consistency parameters as the PVL-delta, but a recency parameter ( 0 < *A* < 1) is used for value updating. The recency parameter indicates how much the expected values of all decks are discounted on each trial.

The PVL-delta is nested within the Value-Plus-Perseverance (VPP) model, which is a hybrid model of the PVL-delta and a heuristic strategy of perseverance. The perseverance decay rate (0 < *k* < 1) decays the perseverance strength of all choices on each trial, akin to how the PVL-decay’s recency parameter affects expected value. The impact of gain ( −∞ < *ε*_*p*_ < ∞) and loss ( −∞ < *ε*_*n*_ <∞) on perseverance parameters reflect how the perseverance value changes after wins and losses, respectively; positive values reflect a tendency to make the same choice, and negative values a tendency to switch choices. The reinforcement learning weight ( 0 < *ω* < 1) is a mixing parameter that controls how much decision-weight is given to the reinforcement learning versus the perseverance term. High and low values reflect more and less reliance on the reinforcement learning term, respectively.

### 2.3. The Orthogonalized Go/NoGo task

Animals use Pavlovian and instrumental controllers when making actions. The Pavlovian controller selects approaching/engaging actions with predictors of appetitive outcomes and avoiding/inhibiting actions with predictors of aversive outcomes. The instrumental controller, on the other hand, selects actions based on the action-outcome contingencies of the environment. The two controllers typically cooperate but sometimes compete with each other (e.g., Dayan, Niv, Seymour, & D Daw, 2006). The orthogonalized Go/NoGo (GNG) task (Guitart-Masip et al., 2012) is designed to examine the interaction between the two controllers by orthogonalizing the action requirement (Go vs. NoGo) against the valence of the outcome (winning vs. avoiding losing money).

Each trial of the orthogonal GNG task has three events in the following sequence: cue presentation, target detection, and outcome presentation. First, one of four cues is presented (“Go to win”, “Go to avoid (losing)”, “NoGo to win”, or “NoGo to avoid”). After some delay, a target (“circle”) is presented on the screen and subjects need to respond with either a *Go* (press a button) or *NoGo* (withhold the button press). Then, subjects receive a probabilistic (e.g., 80%) outcome. See Guitart-Masip et al. (2012) for more details of the experimental design.

The orthogonalized GNG task has been used to study decision-making in healthy populations (Cavanagh et al., 2013), age-related changes in midbrain structural integrity in older adults (Chowdhury, Guitart-Masip, Lambert, Dolan, & Duzel, 2013), and negative symptoms of schizophrenia (Albrecht, Waltz, Cavanagh, Frank, & Gold, 2016). The interaction between Pavlovian and instrumental controllers might also play a role in addiction problems (Guitart-Masip, Duzel, Dolan, & Dayan, 2014). The hBayesDM package currently contains four different models for the orthogonalized GNG task:

1. `gng_m1` (M1 in Guitart-Masip et al., 2012)
  Lapse rate ( 0 < *ξ* <1)
  Learning rate ( 0 < *ε* <1)
  Effective size of a reinforcement ( 0 < *ρ* < ∞)
2. `gng_m2` (M2 in Guitart-Masip et al., 2012)
  Lapse rate ( 0 < *ξ* <1)
  Learning rate ( 0 < *ε* <1)
  Go Bias ( −∞ < *b* < ∞)
  Effective size of a reinforcement ( 0 < *ρ* < ∞)
3. `gng_m3` (M3 in Guitart-Masip et al., 2012)
  Lapse rate ( 0 < *ξ* <1)
  Learning rate ( 0 < *ε* <1)
  Go Bias (−∞ < *b* < ∞)
  Pavlovian bias (−∞ < *π* < ∞)
  Effective size of a reinforcement ( 0 < *ρ* < ∞)
4. `gng_m4` (M5 in Cavanagh et al., 2013)
  Lapse rate ( 0 < *ξ* <1)
  Learning rate ( 0 < *ε* <1)
  Go Bias ( −∞ < *b* < ∞)
  Pavlovian bias ( −∞ < *π* < ∞)
  Effective size of reward reinforcement ( 0 < *ρ*_*rew*_ < ∞)
  Effective size of punishment reinforcement ( 0 < *ρ*_*pun*_ < ∞)

### 2.3.1 GNG: Parameter Descriptions

All models for the GNG include a lapse rate parameter ( 0 < *ξ* < 1), a learning rate parameter ( 0 < *ε* < 1; refer to section 2.2.1 for details), and a parameter for the effective size of reinforcement ( 0 < *ρ* < ∞). The lapse rate parameter captures the proportion of random choices made regardless of the strength of their action probabilities. The *ρ* parameter determines the effective size of reinforcement. The `gng_m4` model has separate effective size parameters for reward ( 0 < *ρ*_*rew*_ < ∞) and punishment ( 0 < *ρ*_*pun*_ < ∞), allowing for rewards and punishments to be evaluated differently.

Three GNG models (`gng_m2, gng_m3`, and `gng_m4`) include a go bias parameter ( −∞ < *b* < ∞). Go bias reflects a tendency to respond (*Go*), regardless of action-outcome associations; high and low values for *b* reflect a high and low tendency to make a go (motor) response, respectively.

Two GNG models (`gng_m3` and `gng_m4`) include a Pavlovian bias parameter ( − ∞ < *π* < ∞). Pavlovian bias reflects a tendency to make responses that are Pavlovian congruent: *π* promotes or inhibits *Go* if the expected value of the stimulus is positive (appetitive) or negative (aversive), respectively.

### 2.4. Probabilistic Reversal Learning task

Environments often have higher-order structures such as interdependencies between stimuli, action, and outcomes. In such environments, subjects need to infer and make use of the structures to make optimal decisions. In the probabilistic reversal learning task (PRL), there exists higher-order structure such that the reward distributions of two stimuli are anticorrelated (e.g. if one option has a reward rate of 80%, the other option has a reward rate of (100-80)%, which is 20%). Subjects need to learn the higher-order structure and take it into account to optimize their decision-making and maximize earnings.

In a typical probabilistic reversal learning (PRL) task, two stimuli are presented to a subject. The choice of a ‘correct’ or good stimulus will usually lead to a monetary gain (e.g., 70%) whereas the choice of an ‘incorrect’ or bad stimulus will usually lead to a monetary loss. The reward contingencies will reverse at fixed points (e.g., Murphy, Michael, Robbins, & Sahakian, 2003) or will be triggered by consecutive correct choices (Cools, Clark, Owen, & Robbins, 2002; Hampton et al., 2006).

The PRL task has been widely used to study reversal learning in healthy individuals (Cools et al., 2002; Gläscher et al., 2009; Ouden et al., 2013). The PRL has been also used to study decision-making deficits associated with prefrontal cortex lesions (e.g., Fellows & Farah, 2003; Rolls, Hornak, Wade, & McGrath, 1994) as well as Parkinson’s disease (e.g., Cools, Lewis, Clark, Barker, & Robbins, 2007; Swainson et al., 2000), schizophrenia (e.g., Waltz & Gold, 2007), and cocaine dependence (Ersche, Roiser, Robbins, & Sahakian, 2008). The hBayesDM package currently contains three models for probabilistic reversal learning tasks:

1. `prl_ewa` (Ouden et al., 2013)
  1 - Learning rate ( 0 < *ϕ* < 1)
  Experience decay ( 0 < *ρ* < 1)
  Inverse temperature ( 0 < *β* < 10)
2. `prl_fictitious` (Gläscher et al., 2009)
  Learning rate ( 0 < *η* < 1)
  Indecision point ( 0 < *α* < 1)
  Inverse temperature ( 0 < *β* < 5)
3. `prl_rp` (Ouden et al., 2013)
  Reward learning rate (0 < *A*_*rew*_ < 1)
  Punishment learning rate ( 0 < *A*_*pun*_ < 1)
  Inverse temperature ( 0 < *β* < 10)

### 2.4.1 PRL: Parameter Descriptions

All PRL models above contain learning rate parameters (refer to section 2.2.1 for details). The `prl_rp` model has separate learning rates for rewards ( 0 < *A_rew_* < 1) and punishments ( 0 < *A_pun_* < 1). In the `prl_ewa` model (Camerer & Ho, 1999), low and high values of *φ* reflect more weight on recent and past outcomes, respectively. All PRL models also contain an inverse temperature parameter (refer to section 2.1.1 for details).

The `prl_ewa` model used in Ouden et al. (2013) contains the decay rate parameter ( 0 < *ρ* < 1). The experienced weight of the chosen option is decayed in proportion to *ρ* and 1 is added to the weight on each trial. Thus, a higher value of *ρ* indicates slower decaying or updating of experienced weight.

The `prl_fictitious` model contains an indecision point parameter ( 0 < *α* < 1). The indecision point reflects a subject’s amount of bias or preference toward an option. High and low values for *α* indicate that there is a greater or smaller preference of one option over the other.

### 2.5. Risk Aversion task

The Risk Aversion (Sokol-Hessner, Camerer, & Phelps, 2012; RA; Sokol-Hessner et al., 2009) task is a description-based task (Hertwig et al., 2004) where possible outcomes of all options and their probabilities are provided to subjects on each trial. In the RA task, subjects choose either a sure option with a guaranteed amount or a risky option (i.e., gamble) with possible gains and/or loss amounts. Subjects are asked to choose which option they prefer (or whether they want to accept the gamble) on each trial. In the RA task, subjects performed two cognitive regulation (*Attend* and *Regulate*) conditions in a within-subject design: in the Attend condition, subjects are asked to focus on each choice in isolation whereas in the Regulate condition, subjects are asked to emphasize choices in their greater context (see Sokol-Hessner et al., 2009 for the details). Data published in Sokol-Hessner et al. (2009) are available in the following paths (paths are also available in RA model help files):

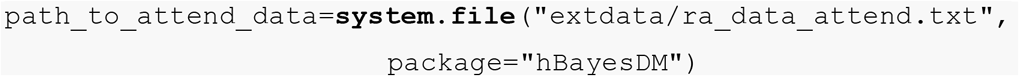

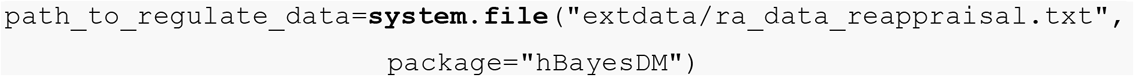

The hBayesDM package currently contains three models for the RA and similar (e.g., Tom, Fox, Trepel, & Poldrack, 2007) tasks:

1. `ra_prospect` (Sokol-Hessner et al., 2009)
  Loss aversion ( 0 < *λ* < 5)
  Risk aversion ( 0 < *ρ* < 2)
  Inverse temperature ( 0 < *τ* < ∞)
2. `ra_noLA` (no loss aversion (LA) parameter; for tasks that involve only gains)
  Risk aversion ( 0 < *ρ* < 2)
  Inverse temperature ( 0 < *τ* < ∞)
3. `ra_noRA` (no risk aversion (RA) parameter; e.g., Tom et al., 2007)
  Loss aversion ( 0 < *λ* < 5)
  Inverse temperature ( 0 < *τ* < ∞)

### 2.5.1 RA: Parameter Descriptions

The `ra_prospect` model contains a loss aversion parameter ( 0 < *λ* < 5), a risk aversion parameter ( 0 < *ρ* < 2) and an inverse temperature parameter ( 0 < *τ* < ∞). See section 2.1.1 for inverse temperature. The risk aversion and loss aversion parameters in the RA models are similar to those in the IGT models. However in RA models, they control the valuation of possible choices under consideration as opposed to the evaluation of outcomes after they are experienced (Rangel et al., 2008).

The `ra_noLA` and `ra_noRA` models are nested within the `ra_prospect` model, where they set loss aversion (`ra_noLA`) or risk aversion (`ra_noRA`) to 1.

### 2.6. Two-Armed Bandit task

Multi-armed bandit tasks or problems typically refer to situations in which gamblers decide which gamble or slot machine to play to maximize the long-term gain. Many reinforcement learning tasks and experience-based (Hertwig, Barron, Weber, & Erev, 2004) tasks can be classified as bandit problems. In a typical two-armed bandit task, subjects are presented with two options (stimuli) on each trial. Feedback is given after a stimulus is chosen. Subjects are asked to maximize positive feedback as they make choices, and they are expected to learn stimulus-outcome contingencies from trial-by-trial experience. The hBayesDM package currently contains a simple model for a two-armed bandit task:

1. `bandit2arm_delta` (Hertwig et al., 2004)
  Learning rate ( 0 < *A* 1)
  Inverse temperature ( 0 < *τ* < 5)

#### 2.6.1 Two-Armed Bandit: Parameter Descriptions

The `bandit2arm_delta` uses the Rescorla-Wagner rule (see Section 2.2.1) for updating the expected value of the chosen option and the softmax choice rule with an inverse temperature (see Section 2.1.1).

### 2.7. The Ultimatum Game (Norm-Training)

The ability to understand social norms of an environment and to adaptively cope with norms is critical for normal social functioning (Gu et al., 2015; Montague & Lohrenz, 2007). The Ultimatum Game (UG) is a widely used social decision-making task that examines how individuals respond to deviations from social norms and adapt to norms in a changing environment.

The UG involves two players: a Proposer and a Responder. On each trial, the Proposer is given some amount of money to divide up amongst the two players. After deciding how to divide the money, an offer is made to the Responder. The Responder can either accept the offer (money is split as offered), or reject it (both players receive nothing). Previous studies show that the most common offer is approximately 50% of the total amount, and “unfair” offers (< ~20% of the total amount) are often rejected even though it is optimal to accept any offer (Güth, Schmittberger, & Schwarze, 1982; Sanfey, 2003; Thaler, 1988). A recent study examined the computational substrates of norm adjustment by using a norm-training UG where subjects played the role of Responder in a norm-changing environment (Xiang et al., 2013).

The UG has been used to investigate social decision-making of individuals with ventromedial prefrontal lesions (Gu et al., 2015; Koenigs et al., 2007) and insular cortex (Gu et al., 2015) lesions, and of patients with schizophrenia (Agay, Kron, Carmel, Mendlovic, & Levkovitz, 2008; Csukly, Polgár, Tombor, Réthelyi, & Kéri, 2011). The hBayesDM package currently contains two models for the UG (or norm-training UG) where subjects play the role of Responder:

1. `ug_bayes` (Xiang et al., 2013)
  Envy ( 0 < *α* < 20)
  Guilt ( 0 < *β* < 10)
  Inverse temperature ( 0 < *τ* < 10)
2. `ug_delta` (Gu et al., 2015)
  Envy (0 < *<* < 20)
  Inverse temperature ( 0 < *τ* < 10)
  Norm adaptation rate ( 0 < *ε* < 1)

#### 2.7.1 UG: Parameter Descriptions

The `ug_bayes` model assumes that the subject (Responder) behaves like a *Bayesian Ideal Observer* (Knill & Pouget, 2004), where the expected offer made by the proposer is updated in a Bayesian fashion. This is in contrast to the `ug_delta` model, which assumes that the subject (Responder) updates the expected offer using a Rescorla-Wagner (delta) updating rule. Both the `ug_bayes` and `ug_delta` models contain envy ( 0 < *α* < 20) and inverse temperature ( 0 < *τ* < 10; refer to section 2.1.1 for details) parameters. The envy parameter reflects sensitivity to the norm prediction error (see below for `ug_bayes` model), where higher and lower values indicate greater and lesser sensitivity, respectively. In the UG, prediction error reflects the difference between the expected and received offer.

In the `ug_bayes` model, the utility of an offer is adjusted by two norm prediction errors: 1) negative prediction errors multiplied by an envy parameter ( 0 < *α* < 20), and 2) positive prediction errors multiplied by a guilt parameter ( 0 < *β* < 20). Higher and lower values for envy (*α*) and guilt (*β*) reflect greater and lesser sensitivity to negative and positive norm prediction errors, respectively. The `ug_delta` model contains only the envy parameter (Gu et al., 2015).

## 3. Mathematical formulation of hierarchical Bayesian models

We first briefly describe HBA (Section 3.1) for readers interested in HBA or a Bayesian framework in general. Then, we illustrate how we program our models using the Stan software package (Carpenter et al., 2016) (**Section 3.2**) and how we formulate hierarchical structures for various types of model parameters (**Section 3.3**). Readers who are not interested in mathematical details may skip **Sections 3.2** and **3.3**.

### 3.1. Hierarchical Bayesian analysis (HBA)

Most computational models do not have closed form solutions and we need to estimate parameter values. Traditionally, parameters are estimated at the individual level with maximum likelihood estimation (MLE): getting point estimates that maximize the likelihood of data for each individual separately (e.g., Myung, 2003). However, individual MLE estimates are often noisy and unreliable especially when there is an insufficient amount of data, which is common in psychology or neuroscience experimental settings (c.f., speeded choice-response time tasks). A group-level analysis (e.g., group-level MLE), which estimates a single set of parameters for the whole group of individuals, may generate more reliable estimates but inevitably ignores individual differences.

For parameter estimation, the hBayesDM package uses HBA, which is a branch of Bayesian statistics. We will briefly explain why hierarchical approaches such as HBA have advantages over traditional MLE methods. In Bayesian statistics, we assume prior beliefs (i.e., prior distributions) for model parameters and update the priors into posterior distributions given the data (e.g., trial-by-trial choices and outcomes) using Bayes’ rule. If Bayesian inference is performed individually for each individual *i*:

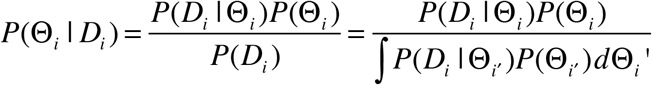

Here, Θ_*i*_ is a set of parameters of a model for individual *i* (e.g., Θ_*i*_ = {*α_i_*, *β_i_*, *γ_i_*, …}, *D_i_* is the data, *p*(*D_i_* | *θ*) is the likelihood (of data given a set of parameters), *P*(*D_i_*) is called evidence (of data being generated by this model), and *P*(Θ*_i_*) and *P*(Θ*_i_* | *D_i_*) are prior and posterior distributions of Θ*_i_*, respectively.

In HBA, hyper-parameters are introduced on top of individual parameters as illustrated in Figure 2A (Gelman, Dunson, & Vehtari, 2013; Kruschke, 2014). If we set hyper-parameters as Φ = {*μ_α_*, *μ_β_*, *μ_γ_*, *σ_α_*, *σ_β_*, *σ_γ_*, …} with group-level normal means *μ*_(.)_ and standard deviations *σ*_(.)_, the joint posterior distribution *P*(Θ,Φ | *D*) is:

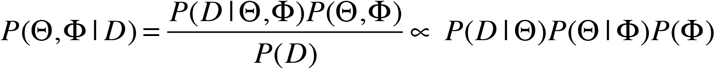

The hierarchical structure of HBA leads to “shrinkage” effects (Gelman et al., 2013) in individual estimates. Shrinkage effects refer to, put simply, when each individual’s estimates inform the group’s estimates, which in turn inform the estimates of all individuals. Consequently, individual parameter estimates tend to be more stable and reliable because commonalities among individuals are captured and informed by the group tendencies (but see Section 6 for its limitations and potential drawbacks). Such a hierarchical approach is particularly useful when the amount of information (e.g., number of trials) from a single person is often too small to precisely estimate parameters at the individual level. A simulation study (Ahn, Krawitz, Kim, Busemeyer, & Brown, 2011a) empirically demonstrated that HBA outperforms individual MLE in parameter recovery (see Figure 2B), which suggests that parameter values estimated with HBA might more accurately reflect individual differences underlying neurocognitive processes than those estimated with individual MLE. Importantly, HBA provides full posterior distributions instead of point estimates, thus it provides rich information about the parameters. HBA also makes it straightforward to make group comparisons in a Bayesian fashion (e.g., comparing clinical and non-clinical groups, see an example in Section 5.4.4). Recent studies in cognitive and decision sciences further confirmed the validity and usefulness of HBA and other hierarchical approaches (e.g., Ahn et al., 2014; Guitart-Masip et al., 2012; Huys et al., 2011; Katahira, 2016; Lee, 2011; Raja Beharelle, Polania, Hare, & Ruff, 2015; Shiffrin, Lee, Kim, & Wagenmakers, 2008).

**Figure 2.**
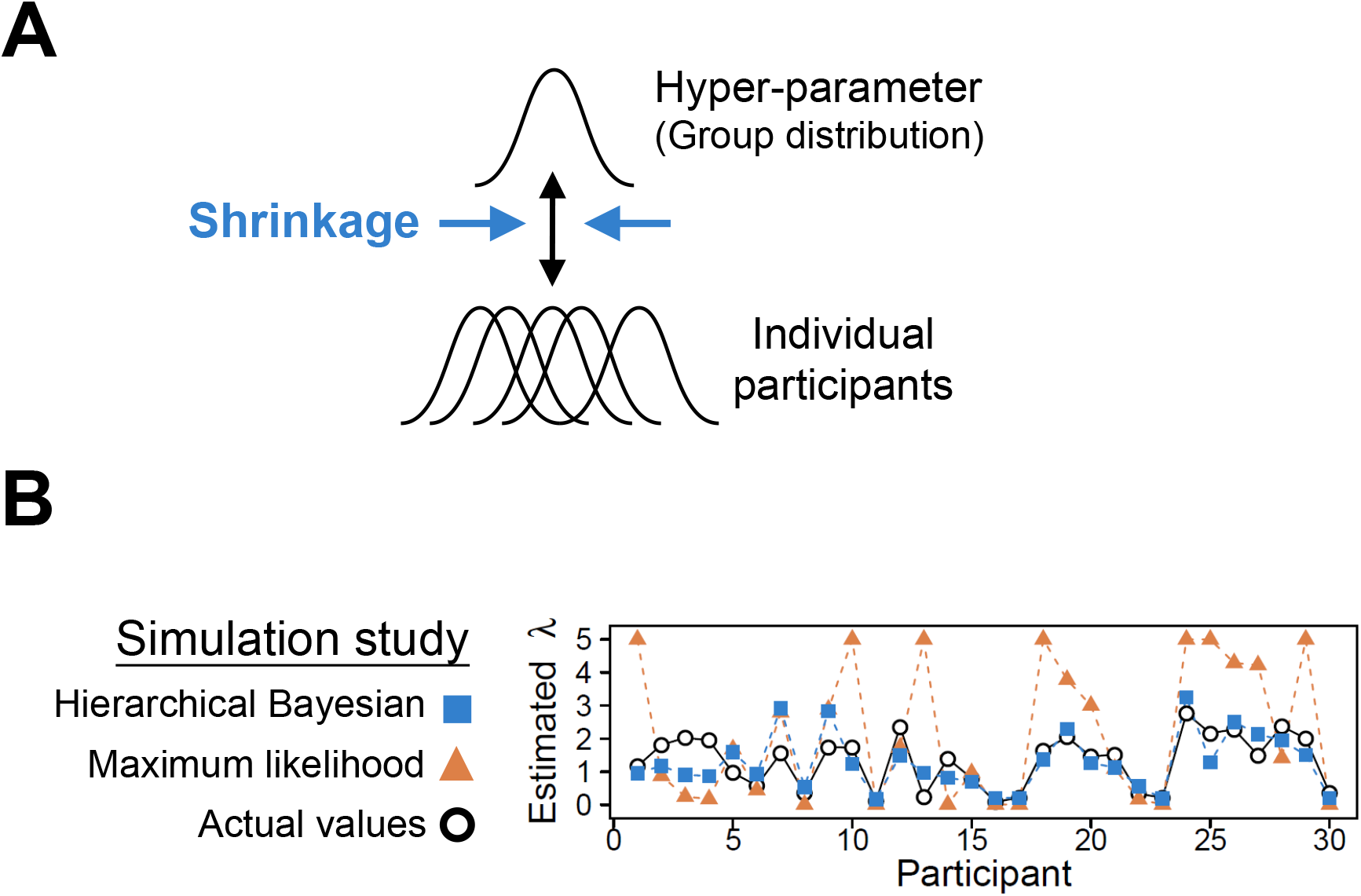
**(A)** A schematic illustration of hierarchical Bayesian analysis (HBA). In this example, individual parameters are assumed to come from a hyper (group) parameter. **(B)** The results of a parameter recovery study (Ahn et al., 2011) between HBA and maximum likelihood estimation (MLE). Thirty subjects’ data on the IGT were simulated from true parameters (black circles) and parameters were estimated with hierarchical Bayesian analysis (blue squares = individual posterior means) and individual maximum likelihood estimation (red triangles). The performance of the two approaches is shown for the loss aversion parameter (*λ*).

## 4. Performing hierarchical Bayesian analysis with Stan

In the hBayesDM package, posterior inference for all models is performed with a Markov Chain Monte Carlo (MCMC) sampling scheme using a newly developed probabilistic programming language Stan (Carpenter et al., 2016) and its R instantiation RStan (http://mc-stan.org/interfaces/rstan). Stan uses a specific MCMC sampler called Hamiltonian Monte Carlo (HMC) to perform sampling from the posterior distribution. During each iteration of HMC, derivatives of the density function, together with the auto-optimized Metropolis acceptance rate and step size and maximum steps, are utilized to find out the direction of the target posterior distribution (Carpenter et al., 2016). HMC offers more efficient sampling than conventional algorithms implemented in other software like BUGS (Lunn, Thomas, Best, & Spiegelhalter, 2000; Lunn, Spiegelhalter, Thomas, & Best, 2009) and JAGS (Plummer, 2003). Moreover, HMC works well even for complex models with high-dimensional model structures and highly correlated model parameters. A drawback of HMC is that it is not capable to directly sample discrete parameters because HMC uses derivatives of the density. Yet one could marginalize the posterior density to obtain discrete outcomes. See the Stan reference manual (http://mc-stan.org/documentation/) and Kruschke (2014; Chapter 14) for a comprehensive description of HMC and Stan. To learn more about the basic foundations of MCMC, see Krushcke (2014; Chapter 7).

To use the hBayesDM package, users do not need to know how to program in Stan. However, for those interested in understanding our models and Stan in general, we briefly introduce the general structure of model specification in Stan, followed by a detailed hierarchical parameter declaration and optimizing approaches that are utilized in hBayesDM. Lastly, we describe how we calculate log likelihood and model-fits inside Stan models.

### 4.1. General structure of Stan model specification

Many useful features of BUGS were incorporated into Stan’s design; thus, Stan is similar to BUGS (or JAGS) and users who are familiar with BUGS would find Stan relatively easy to use (see Stan reference manual, Appendix B: http://mc-stan.org/documentation/). There are six model blocks in the general structure of Stan model specification, as listed below. Note that Stan implements sequential execution in its model specification unlike BUGS and JAGS where the order of code does not affect a model’s execution:

~~~
data {
… read in external data…
}
transformed data {
… pre-processing of data …
}
parameters {
… parameters to be sampled by HMC …
}
transformed parameters {
… pre-processing of parameters …
}
model {
… statistical/cognitive model …
}
generated quantities {
… post-processing of the model …
}
~~~

Note that the data, parameters and model blocks are mandatory in Stan, whereas transformed data, transformed parameters and generated quantities blocks are optional. Nonetheless, we typically use all these optional blocks in hBayesDM for different purposes: (1) We use the transformed data block to maintain a concise programming style and assign initial values. (2) We implement non-centered parameterization (a.k.a., Matt Trick) in the transformed parameters block to optimize sampling and reduce the autocorrelation between group-level parameters in our hierarchical models. Details will be explained in the optimizing section of the tutorial (Section 4.3). (3) We include the generated quantities section to explicitly calculate log-likelihood of the corresponding model and compute out-of-sample prediction accuracy (see Section 4.4) for model comparison.

### 4.2. Hierarchical parameter declaration in Stan

When declaring hierarchical parameters in hBayesDM with Stan, we assume that individual parameters are drawn from group-level normal distributions. Normal and half-Cauchy distributions are used for the priors of the group-level normal means (*μ*_(.)_) and standard deviations (*σ*_(.)_), respectively. We employ flat (uniform) or weakly informative priors (Gelman et al., 2013) to minimize the influence of those priors on the posterior distributions when sample sizes are small. We used standard normal priors for group-level means (e.g., Lee, 2011; Shiffrin et al., 2008; Wetzels, Vandekerckhove, & Tuerlinckx, 2010), which also makes it easy to optimize Stan codes (see Section 4.3). For the group-level standard deviations, we used half-Cauchy prior distributions, which tend to give sharper and more reasonable estimates than uniform or inverse-Gaussian prior distributions (Gelman, 2006). According to the range of parameters of interest, we introduce four ways of declaring hierarchical parameters: unbounded parameters, positive parameters, parameters bounded between 0 and 1, and parameters bounded between 0 and an upper limit *U*.

For unbounded parameters (say *ξ* for a general individual parameter for illustration purposes), we declare:

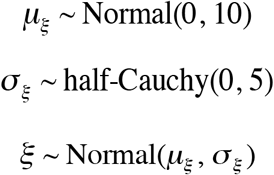

where *μ_ξ_* (group mean parameter) is drawn from a wide normal distribution, *σ_ξ_* (group standard deviation parameter) is drawn from a positive half-Cauchy distribution, and *ξ* is distributed as a normal distribution with a mean of *μ_ξ_* and a standard deviation of *σ_ξ_*. Note that we use the wide normal distribution (weakly-informative prior) to keep the prior bias minimum. Plus, the use of the positive-half Cauchy ensures most density is between 0 and 10, meanwhile the HMC sampler is still able to visit beyond its upper bound, resulting in a soft constrain (Gelman et al., 2013).

For positive parameters (e.g., the effective size of reinforcements in the orthogonalized GNG task), we apply an exponential transformation to constrain an unbounded parameter to be greater than 0, such that the transformed prior is exclusively positive and avoids extreme values. Note that it results in a small bias toward zero. In hBayesDM, we define:

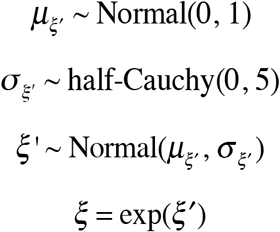

For parameters that are bounded between 0 and 1 (e.g., learning rate), we use the inverse probit transformation (the cumulative distribution function of a unit normal distribution) to convert unconstrained values into this range. In fact, given the mathematical relationship between the probability density function (pdf) and the cumulative density function (cdf) of the unit normal distribution, this transformation guarantees the converted prior to be uniformly distributed between 0 and 1. Several studies have demonstrated the robustness and effectiveness of this transformation (e.g., Ahn et al., 2014; Wetzels et al., 2010). To effectively implement this, Stan provides a fast approximation of the inverse probit transformation (i.e., `Phi_approx` function), which we adopted:

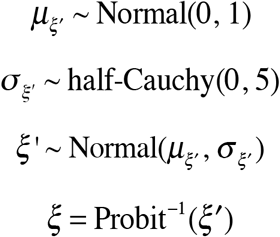

For parameters that are bounded between 0 and an upper limit *U* (e.g., inverse Softmax temperature, loss aversion in the Risk-Aversion task), we simply adapt the declaration rule for [0, 1] parameters and multiply it by the upper limit *U*. Likewise, the converted prior is distributed as a uniform distribution between 0 and *U*. If *U* is relatively small (less than ˜20) and, we used this approach instead of using a positive parameter (with an exponential transformation) to keep the prior bias minimum. When we used such an upper bound, posterior fits are checked to ensure that the parameter estimates are not very close to the boundary. Formally, we declare:

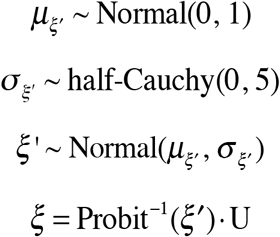

As shown above, we do not employ truncated sampling in declaring hierarchical parameters because hard constrain (e.g., *ξ*~ Normal(0, 1)T[0, *U*]) may harm the HMC sampling algorithm and return poorly converged posterior distributions (Carpenter et al., 2016). If users want to build their own hierarchical Bayesian models for their research, they can refer to our practice of standardizing parameter declaration.

### 4.3. Optimizing approaches in Stan

Hierarchical models often suffer from highly correlated group-level parameters in posterior distributions, creating challenges in terms of model convergence and estimation time (Gelman et al., 2013; Kruschke, 2014). In order to address the challenges, we practice reparameterization and vectorization to optimize the model specification in hBayesDM.

A Normal(*µ*, *σ*) distribution, like other distributions in the location-scale distribution family, can be reparameterized to be sampled from a unit normal distribution that is multiplied by the scale parameter *σ* and then shifted with the location parameter *µ*. Formally,

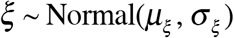

is mathematically equivalent to

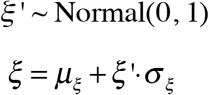

Such transformation is referred to as the non-centered parameterization (a.k.a., Matt Trick) by the Stan Development Team (2016), and it effectively reduces the dependence between *μ_ξ_*, *ξ*, and *σ_ξ_* and increases effective sample size.

In addition to reparameterization, we use vectorization to improve MCMC sampling. For example, suppose one experiment consists of N participants, then its individual level parameter *ξ* is an N-dimensional vector. Instead of declaring *ξ* as:

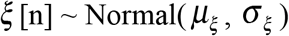

we vectorize it as:

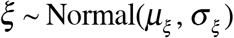

to make full use of Stan’s vectorization of all sampling statements. As a rule of thumb, one may want to use vectorization as long as it is possible. All hBayesDM’s models that implement both reparameterization and vectorization can be found under the directory of `…\R\R-x.x.x\library\hBayesDM\stan`, or the path can be retrieved by calling the following R command: `file.path(.libPaths()`, “`hBayesDM`”, “`stan`”). Those interested in more details of optimizing Stan models can read Stan reference manual (http://mc-stan.org/documentation/, “Optimizing Stan Code” Chapter).

### 4.4. Computing log-likelihood inside Stan models

The hBayesDM package provides two model performance indices including Leave-One-Out Information Criterion (LOOIC) and Widely Applicable Information Criterion (WAIC). We follow Vehtari et al. (2016) to compute and monitor Stan’s pointwise log-likelihood in the generated quantities block. The generated quantities block serves as the post-processing of the model, with its commands being executed only after the HMC sampling. Therefore, it does not significantly increase the time required for Bayesian inference. The generated quantities block is particularly useful when users intend to monitor pointwise log-likelihood (Vehtari, Gelman, & Gabry, 2016), reproduce predictive values or obtain internal model variables. Practically, we initialize the pointwise log-likelihood to be 0 for each participant, then we repeat the same model of the ‘model’ block in the generated quantities block, except we replace the sampling statement with the explicit computation of pointwise log-likelihood. Please be aware that in many RLDM tasks (especially RL tasks), choices on one trial are dependent on those on other trials. Therefore, instead of gathering the trial-by-trial log-likelihood, we sum them over per participant and obtain the pointwise log-likelihood at the participant level. Below is the pseudocode as an example of what is described above:

~~~
model {
…
 for (i in 1:N) {
 for (t in 1:T) {
  Choice[i, t] ~ categorical_logit(ev);
…
}
Generated quantities {
…
 for (i in 1:N) {
  log_lik[i]=θ;
  for (t in 1:T) {
   log_lik[i]=log_lik[i] + categorical_logit_lpmf(Choice[i, t] | ev);
…
}
~~~

Once having the pointwise log-likelihood per participant, it is straightforward to compute LOOIC and WAIC (Vehtari et al., 2016). Both LOOIC and WAIC provide the estimate of out-of-sample predictive accuracy in a fully Bayesian way, which samples new participants from the hierarchical group, generates new data from those new participants and evaluates how well a model make predictions on the new data set. What makes LOOIC and WAIC more reliable compared to Akaike information criterion (AIC; Akaike, 1987; Bozdogan, 1987) and deviance information criterion (DIC; Spiegelhalter, Best, Carlin, & van der Linde, 2002) is that both LOOIC and WAIC use the pointwise log-likelihood of the full Bayesian posterior distribution, whereas AIC and DIC are based only on point estimates to calculate model evidence. We used functions included in the loo package (Vehtari et al., 2016) to generate LOOIC and WAIC values. Both LOOIC and WAIC are on the information criterion scale; thus, lower values of LOOIC or WAIC indicate better out-of-sample prediction accuracy of the candidate model.

## 5. Step-by-step tutorials of the hBayesDM package

### 5.1. Installing *hBayesDM*: Prerequisites

Before installing hBayesDM, it is necessary to have an up to date version of R (version 3.2.0 or later is recommended) and RStan on your machine. RStudio (www.rstudio.com) is not required but strongly recommended. Typically, RStan can be installed just by entering the following command into the R console:

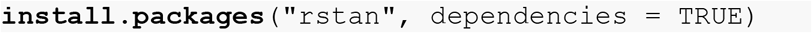

For Windows, it is necessary to install *Rtools* before installing RStan. Instructions for installing Rtools on a Windows machine can be found in this link (https://github.com/stan-dev/rstan/wiki/Install-Rtools-for-Windows).

After RStan (and Rtools for Windows users) is installed, it is recommended to restart R (or RStudio) and test the installation before moving on to install hBayesDM. This can be accomplished by trying to fit the “Eight Schools” example that is provided on RStan’s *Getting Started* page (https://github.com/stan-dev/rstan/wiki/RStan-Getting-Started).

### 5.2. Installing *hBayesDM*

The hBayesDM package is available on the Comprehensive R Archive Network (CRAN). Each software package that is submitted to CRAN undergoes daily checks to ensure that the package is functional and without major bugs. CRAN also makes it very simple to install a package, so this is the preferred method for installation. To install hBayesDM from CRAN, use the following call:

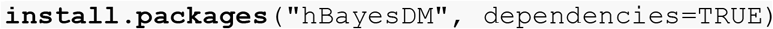

### 5.3. How to use hBayesDM: Navigating

After hBayesDM has been installed correctly, the package must be loaded into the current environment. Users will be able to access all the functions that are included with the package after hBayesDM is loaded. To load hBayesDM, use the following command:

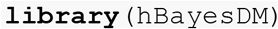

After loading the package, users should see a message that displays the version number of the current hBayesDM install. For a list of all the models available in the package, one can refer to the package help files with the following command:

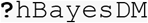

This will bring up a help file that contains a general description of the package along with a list of all RLDM tasks and models that can be fit with hBayesDM. One can follow the links provided in the help file to find in-depth documentation describing how to fit each model.

### 5.4. How to use hBayesDM: Model Fitting

The conceptual framework of computational modeling and four steps of doing HBA with hBayesDM are illustrated graphically in Figure 3. These steps are described in further detail below. To exemplify these steps, four models of the orthogonalized GNG task are fit and compared using the hBayesDM package. As a reminder – users can refer to the help file of any model to learn how to run a real-data example. Also, commands and input arguments for running or evaluating a model are very similar or the same for all models. Thus, if users learn how to run one model, they can also easily run other models.

**Figure 3.**
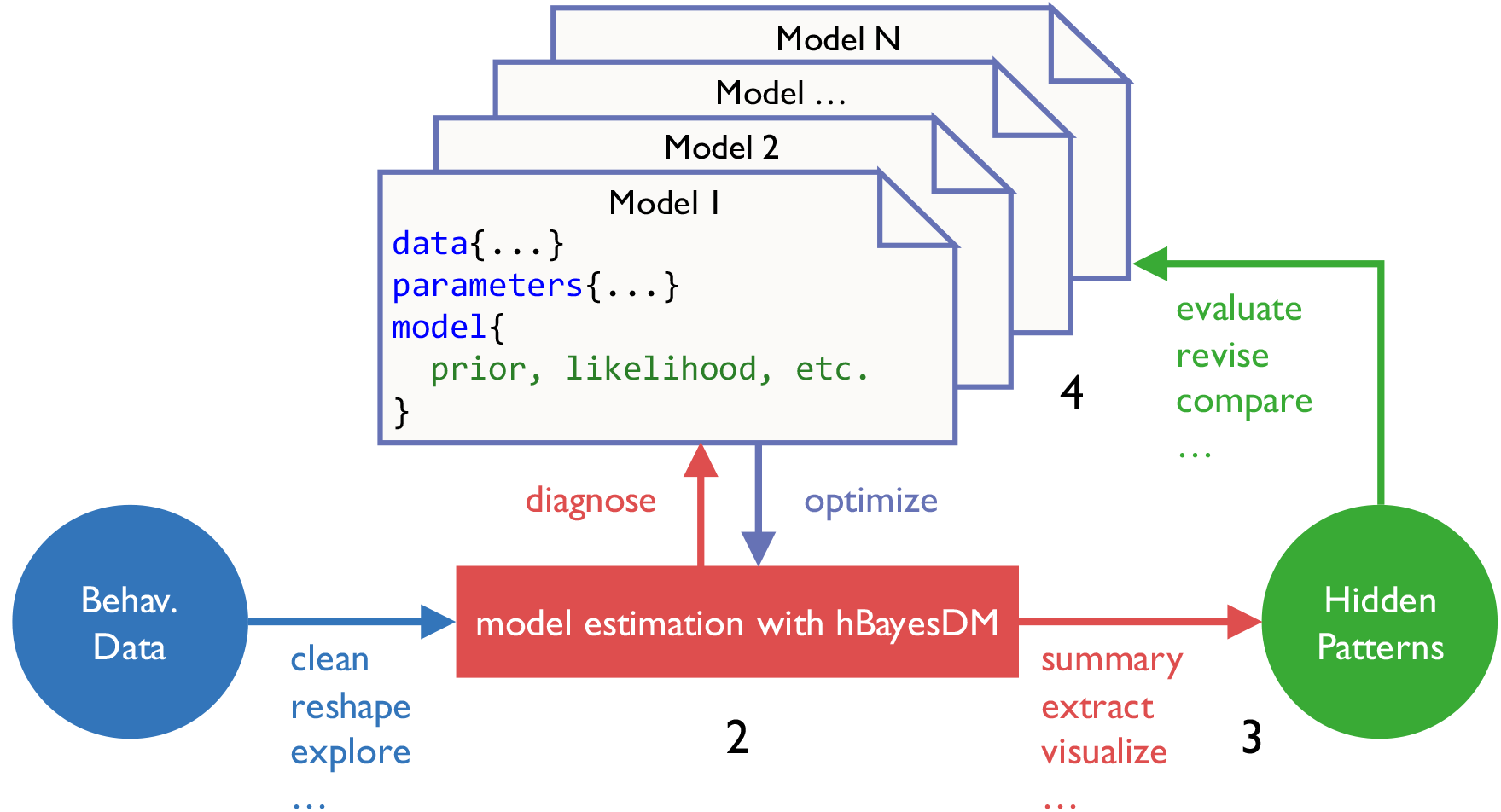
Pipeline of performing computational modeling with hBayesDM. There are 4 steps to perform hierarchical Bayesian analysis (HBA), (1) prepare data, (2) fit candidate models, (3) extract and visualize parameters and/or variables, and (4) model comparison (see details in the text).

#### 5.4.1. Prepare the data

To begin, all subjects’ data (for the current analysis) should be combined into a single text (*.txt) file where rows represent trial-by-trial observations and columns represent variables of interest. The first row of the text file must contain the column headers (names) of the variables of interest.

Subjects’ data must contain variable headers that are consistent with the column names specified in the model help file (see **Table 1**). For example, in the orthogonalized GNG task, there should exist columns labeled: “subjID”, “cue”, “keyPressed”, and “outcome”, where subjID is a subject-specific identifier, cue is a nominal integer specifying the cue shown on the given trial, keyPressed is a binary value representing whether or not a key was (1) or was not (0) pressed on the given trial, and outcome represents a positive (1), negative (-1), or neutral (0) outcome on the given trial. The text file may also contain other data/column headers, but only the aforementioned variables will be used for modeling analysis. All of the above information for each model can be found in the package help files, which can be accessed with R’s help command (e.g., for orthogonalized GNG model 1: **?**`gng_m1`). Across all the models implemented in hBayesDM, the number of trials within the data file is allowed to vary across subjects, but there should exist no N/A data. If some trials contain N/A data (e.g., `outcome=NA`), remove these trials before continuing. If trials containing N/A data are not removed prior to model fitting, they will be removed automatically and the user will receive a warning message.

Sample data can be retrieved from the package folder with the R command shown below. Note that the file name of sample (example) data for a given task is **taskName_exampleData.txt** (e.g., dd_exampleData.txt, igt_exampleData.txt, gng_exampleData.txt, etc.):

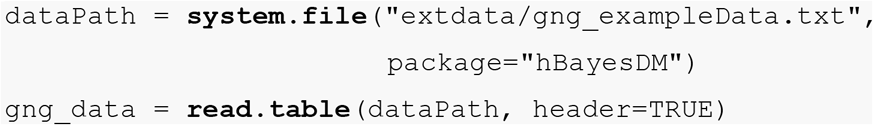

If data are downloaded from an external source to “/home/user1/Downloads”, the user may specify the path using a character string like below:

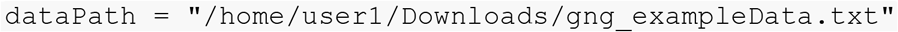

#### 5.4.2. Fit candidate models

Since hBayesDM uses MCMC sampling to generate posterior distributions, there are many arguments that may be passed to Stan through the model functions in order to fine-tune the sampling behavior. There are also arguments that can be used for user convenience. **Table 2** shows arguments that are common to all model functions in hBayesDM. Note that in the table an asterisk (*) denotes an argument that may unpredictably change the computation time and/or sampling behavior of the MCMC chains (Homan & Gelman, 2014). For this reason, it is advised that only advanced users alter the default values of these arguments.

Below, the `gng_m1` model is fit using the sample data that comes with the package. The command indicates that three MCMC chains are to be run and three cores are to be used for parallel computing. Note that parallel computing is only useful for multiple chains; it is common to use one core per chain to maximize sampling efficiency. If “`example`” is entered as an argument for `data`, hBayesDM will use the sample data for the task. Convenience arguments such as `saveDir` can be used in order to save the resulting model output to a local directory. This is useful for when model fitting is expected to take long periods of time and users want to ensure that the data are saved. Also, the email argument allows users to be notified by an email message upon the completion of model fitting.

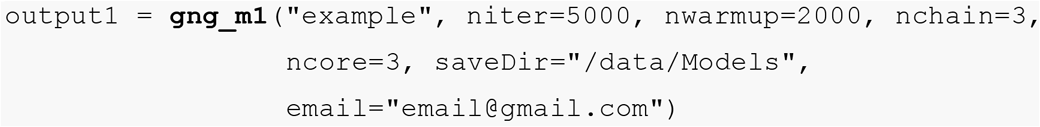

A model function has default values for all arguments except for `data`, so the above command is equivalent (aside from `saveDir` and `email` arguments) to the more concise call below:

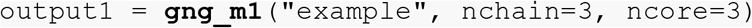

If the `data` argument is left blank, a file browser window will appear, allowing the user to manually select the text file with their file browser. The default input arguments for each model were selected based on our experience with the sampling behavior of each model with respect to the data we have access to. For each model being fitted, `niter` and `nwarmup` values (and control parameters for advanced users) might need to be experimented with to ensure that convergence to target posterior distributions is being reached. Later sections will discuss convergence in more detail.

Executing any model function command in hBayesDM will generate messages for the user within the R console, exemplified by Figure 4A. It will take approximately up to 3 minutes (with the `gng_m1` model & “`example`” data) for the model fitting to complete. Note that you may get warning messages about “numerical problems” or that there are a certain number of “divergent transitions after warm-up”. When we check our models with example datasets, warning messages appear mostly at the beginning of the warm-up period, and there are very few divergent transitions after warm-up. In such cases, the warnings can be ignored. For a technical description of these (and other) sampling issues, see Appendix D of the Stan Reference Manual. When the model fitting is complete, the R console will print the message in Figure 4B. The output data will be stored in `output1`, a class `hBayesDM` object containing a list with 6 following elements:

1. `model`:
  Name of the fitted model (i.e., `output1$model` is “`gng_m1`”)
2. `allIndPars`:
  Summary of individual subjects’ parameters (default: posterior *mean values of individual parameters*). Users can also choose to use posterior *median* or *mode* in a model function command (e.g., `indPars=`“`mode`”)). See Figure 4C to view the values of `allIndPars` for `gng_m1` printed to the R console.
3. `parVals`:
  Posterior MCMC samples of all parameters. Note that hyper (group) posterior mean parameters are indicated by `mu_PARAMETER` (e.g., `mu_xi, mu_ep, mu_rho`). These values are extracted from `fit` with RStan’s `extract`()function.
4. `fit`:
  An `rstan` object that is the output of RStan’s `stan`() function. If users would like to use Rstan commands, they should be performed on this object. See Figure 4D for a summary of `fit` printed to the R console.
5. `rawdata`:
  Raw trial-by-trial data used for HBA. Raw data are provided in the output to allow users to easily access data and compare trial-by-trial model-based regressors (e.g., prediction errors) with choice data.
6. `modelRegressor` (optional):
  Trial-by-trial model-based regressors such as prediction errors, the value of the chosen option, etc. For each model, we pre-selected appropriate model-based regressors. Users can refer to the package help files for the details. Currently (version 0.2.3.2), this feature is available only for the orthogonalized GNG task.

**Figure 4.**
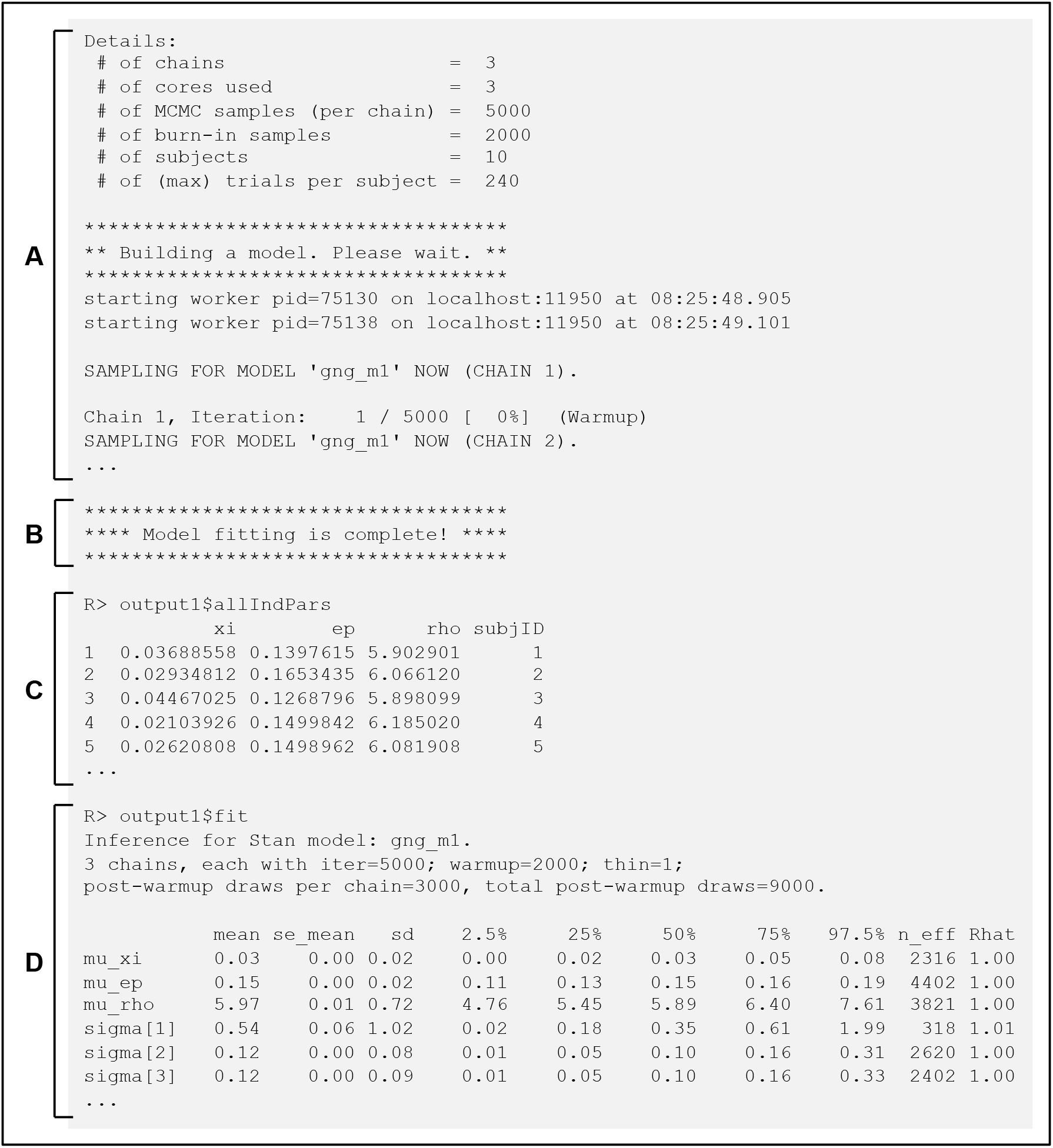
Panel (A) shows the message displayed in the R console after a model function is called. Here, “Details” shows information relevant to both the arguments passed to the function and to the data that was specified by the user. The console also shows the progression of the MCMC sampling. As shown in Panel (B), upon the completion of model fitting, a message is presented to the user. Panel (C) and (D) show that users can retrieve the summary statistics of individual model parameters and Stan model fits (the Stan fit object stored as `output1`).

#### 5.4.3. Plot model parameters

It is important to both visually and quantitatively diagnose MCMC performance (i.e., visually check whether MCMC samples are well mixed and converged to stationary distributions). For the visual diagnostics of hyper (group) parameters, users can call `plot.hBayesDM()` or just `plot()`, which searches for an extension function that contains the class name. The class of any hBayesDM output is `hBayesDM`. For a quantitative check on convergence, the Gelman-Rubin convergence diagnostic (Gelman & Rubin, 1992) for each parameter is computed by RStan and stored in the `fit` element of the hBayesDM model output. To see the Gelman-Rubin values, refer to Figure 4D. Here, *Ȓ* (`Rhat`) is the Gelman-Rubin index used to assess the convergence of the MCMC samples. *Ȓ* values close to 1.00 would indicate that the MCMC chains are converged to stationary target distributions. *Ȓ* values greater than 1.1 are typically considered to represent inadequate convergence. For all models included in hBayesDM, *Ȓ* values are 1.00 for most parameters or at most 1.04 when tested on example datasets.

Users can also use trace plots to visually check MCMC samples. The command shown below (with font size set to 11) shows how to use the `plot()` command to create trace plots of hyper (group) parameters (see Figure 5A for an example):

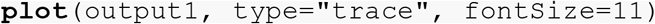

The trace plots indicate that the MCMC samples are indeed well mixed and converged, which is consistent with their *Ȓ* values. Note that the plots in Figure 5A exclude burn-in samples. Users can include burn-in (warm-up) MCMC samples to better understand sampling behavior if necessary. The following function call produces the plot in Figure 5B that includes burn-in samples:

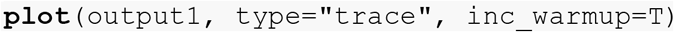

Users can also plot the posterior distributions of the hyper (group) parameters with the default `plot()` function by not specifying the type argument. The following function call produces the plot in Figure 5C.

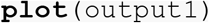

**Figure 5.**
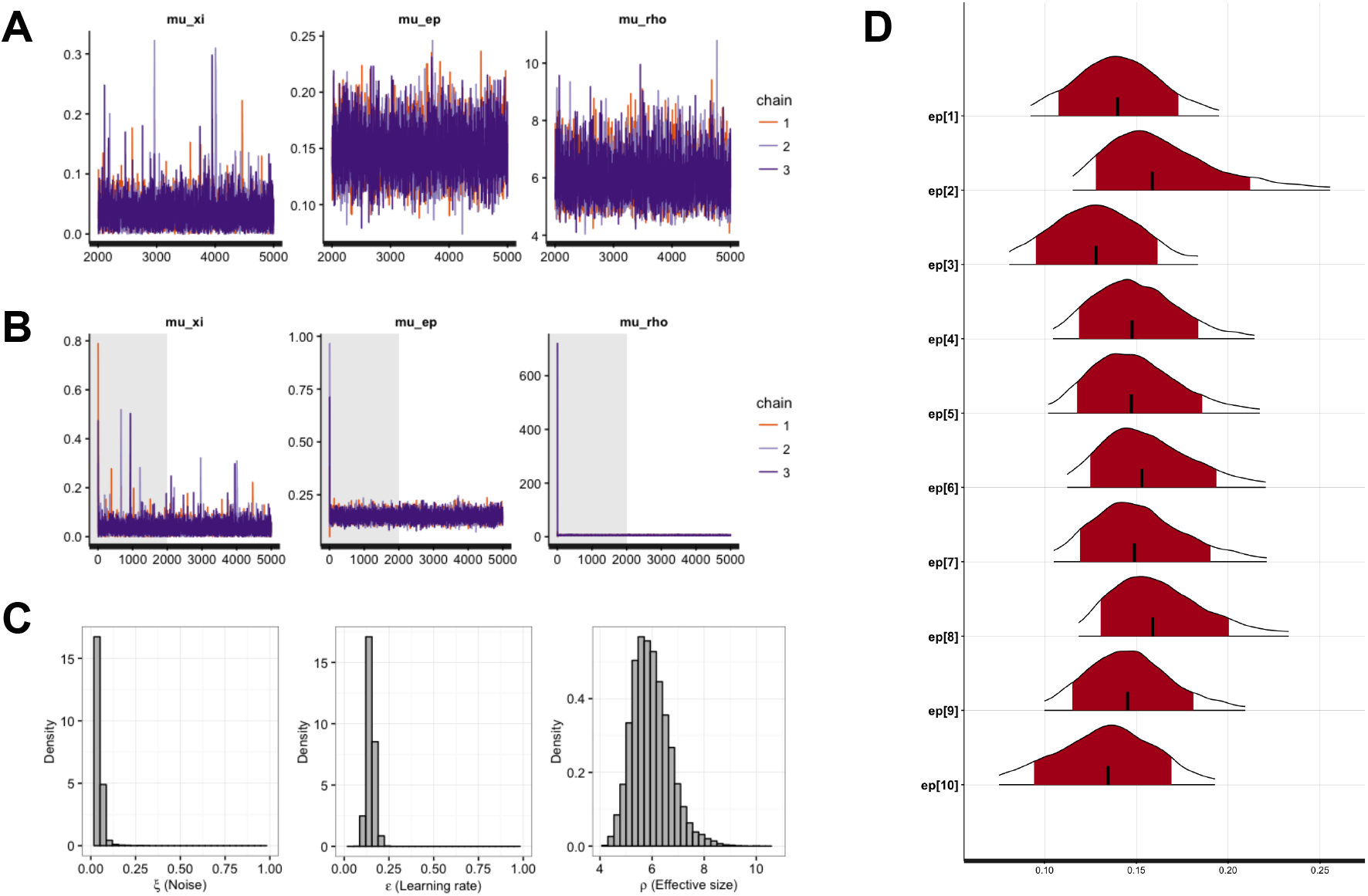
(A) Traceplots for the group-level (hyper) parameters of the `gng_m1` model. The 3 chains show excellent mixing, suggesting that they have converged to their target distributions. (B) The same traceplots as Panel (A), however, these also include the warm-up (burn-in) samples, highlighted by the gray background shading. (C) The posterior distributions of the group-level (hyper) parameters. (D) Individual-level posterior distributions. The red shading and tailed white areas represent the 80% and 95% kernel density estimates, respectively. Note that all plots above are generated directly from hBayesDM and RStan functions, with no further modifications.

To visualize individual parameters, users can use the `plotInd()` command. The following call plots each individual’s *ε* (learning rate) parameter (see Figure 5D):

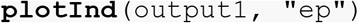

#### 5.4.4. Compare models (and groups)

To compare multiple models using LOOIC or WAIC values, the first step is to fit all models in the same manner as the `gng_m1` example above. The following commands will fit the rest of the orthogonalized Go/Nogo models available within hBayesDM:

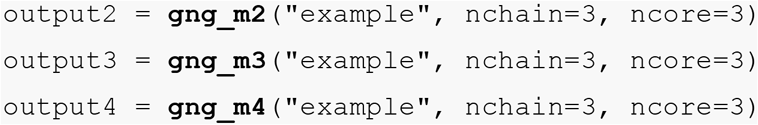

Note that each model should be checked for convergence in the same manner as `gng_m1`. If for any reason a model fails to converge, re-fit the model after model diagnostics (see **5.4.6**) or exclude the model from model comparisons.

Next, users can assess model fits using the `printFit()` command, which is a convenient way to summarize LOOIC and WAIC of all considered models. Assuming all four models’ outputs are named `output1` (gng_m1), `output2` (gng_m2), `output3` (gng_m3), and `output4` (gng_m4), their model fits can be simultaneously summarized by the following command, the results of which are illustrated in Figure 6A:

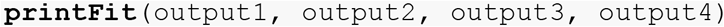

The lower LOOIC or WAIC values indicate better model performance; thus, the model number 4 has the best LOOIC and WAIC compared to other models. Users interested in more detailed information including standard errors and expected log pointwise predictive density (elpd) can use the `extract_ic()` function (e.g., extract_ic(output3)) to extract this information. Note that the `extract_ic()` function can be used only for a single model output, unlike `printFit()`.

**Figure 6.**
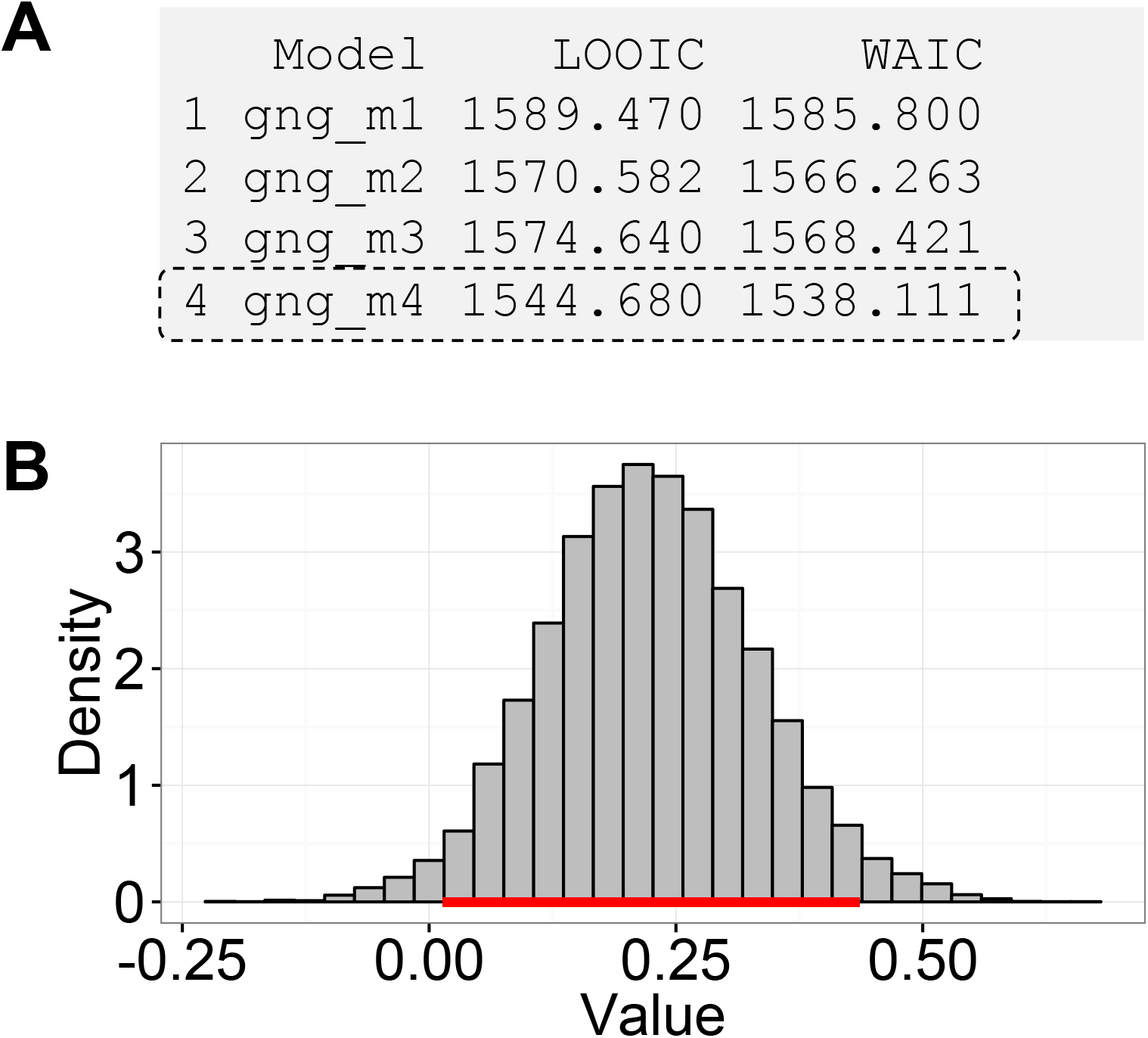
**(A)** An output of the `printFit()` command, which prints model performance indices (LOOIC and WAIC) of competing model(s). The resulting table shows the name of each model, followed by their LOOIC and WAIC values. Lower LOOIC and WAIC values correspond to better model performance. Here, `gng_m4` (highlighted with a dashed box) has the lowest values. **(B)** The result of the `plotHDI()` function that plots the 95% Highest Density Interval (HDI) of the posterior distribution difference between two group parameters. The red bar indicates the 95% HDI.

There exist other model comparison methods including the simulation method (a.k.a., absolute model performance) (Ahn et al., 2008; 2014; Guitart-Masip et al., 2012; Steingroever, Wetzels, & Wagenmakers, 2013), parameter recovery (Ahn et al., 2014; Ahn, Krawitz, Kim, Busemeyer, & Brown, 2011a), and generalization criterion (Ahn et al., 2008; Busemeyer & Wang, 2000). Models that show the best goodness-of-fit may not perform well on other indices (e.g., Ahn et al., 2014), so it is recommended that researchers use multiple model comparison methods if at all possible.

#### 5.4.5. Group comparisons

Having selected the best-fit model, users may want to use the model to compare the parameter estimates of different populations. With a hierarchical Bayesian framework, users can compare model parameters of multiple groups or within-subject conditions in fully Bayesian ways (e.g., Ahn et al., 2014; Chan et al., 2014; Fridberg, Ahn, Kim, Bishara, & Stout, 2010; Kruschke, 2014; Vassileva et al., 2013). The (posterior) distributions show the uncertainty in the estimated parameters and we can use the posterior highest density interval (HDI) to summarize the uncertainty. 95% HDI refers to “the span of values that are most credible and cover 95% of the posterior distribution” (Kruschke, 2014). To examine the difference of a particular parameter between two groups, we can calculate the difference of the hyper-distributions across the groups, and examine its credible interval (i.e., its 95% HDI) (Kruschke, 2010; 2011). Note that this is different from testing a null hypothesis (e.g., test if two groups are the same or not on the parameter of interest), for which Bayesian hypothesis testing (e.g., Bayes factor) (Kass & Raftery, 1995; Myung & Pitt, 1997; Wagenmakers, 2007) or a region of practical equivalence (ROPE) around the null value should be used (Kruschke, 2014; 2011).

As an example, let’s compare two groups’ model parameters in a Bayesian fashion. First, prepare each group’ data as separate text files:

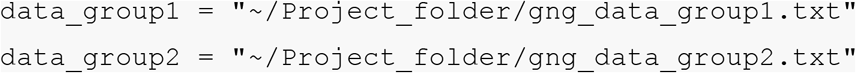

Here, `gng_data_group1.txt` and `gng_data_group2.txt` contain all group 1 subjects’ and group 2 subjects’ data, respectively. Next, the model is fit in the same manner as before on each group separately. We recommend the same number of chains and MCMC samples be used for each group:

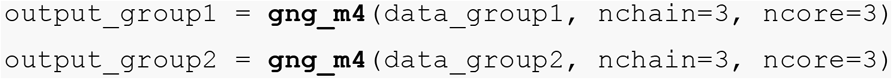

Make sure to check if MCMC samples are well mixed and converged to stationary distributions (Section 5.4.3). Next, compute the difference between the hyper (group) parameters of interest by making a simple subtraction. For example, if we want to compare the Pavlovian bias parameter (*π*) across the two groups:

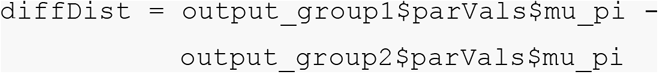

The above command subtracts the `mu_pi` parameter of group 2 from that of group 1. Note that these parameter values are stored within the `parVals` element of an hBayesDM object. To generate the credible interval of the difference between the groups, users can use the following command, which will print the 95% HDI to the R console:

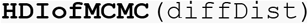

Users can also visually inspect 95% HDI with the following command (95% HDI is also printed to the R console with the command):

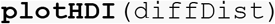

Figure 6B shows the result of the `plotHDI()` command above. The red bar along the bottom of the plot encompasses the 95% HDI.

#### 5.4.6. Improving sampling performance in hBayesDM

When chains fail to reach convergence (e.g., *Ȓ* > 1.10 or MCMC chains are poorly mixed when visually inspected), users are recommended to modify several HMC sampling parameters to improve the performance. Though model performance may be model and parameter specific, we provide a general approach for users to experiment with. Three sampling parameters are relevant for sampling performance: the Metropolis acceptance rate (*δ*, default=0.8), the initial HMC step size (*ε*, default=1.0), and the maximum HMC steps per iteration (*L*; i.e., maximum tree depth, default=10). We refer readers to a Stan help file (`?stan`) for more details. With default sampling parameters and sample datasets, all models implemented in the hBayesDM package showed excellent convergence and mixing of MCMC chains. However, if users notice any signs of poor convergence or mixing, we suggest users increase *δ*, decrease *ε*, and/or increase *L*. The adjustment is hBayesDM is illustrated below (taking gng_m1 as an example):

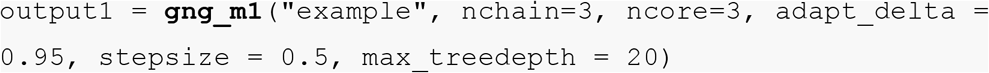

Be aware that such adjustment might dramatically increase the model estimation time and does not necessarily guarantee an improved sampling performance. The failure of an adjusted model estimate might further suggest that such model is not suitable for the current dataset, and one needs to consider using alternative models to fit the data. If users encounter a problem and would like to seek help from the hBayesDM developers, they can ask questions to our mailing list (https://groups.google.com/forum/#!forum/hbayesdm-users).

### 5.5. Extracting trial-by-trial regressors for model-based fMRI/EEG analysis

In model-based fMRI or EEG (Mars et al., 2008; e.g., O'Doherty et al., 2007), model-based time series of a latent cognitive process are generated by computational models, and then time series data are regressed again fMRI or EEG data. This model-based neuroimaging approach has been particularly popular in cognitive neuroscience (e.g., Ahn, Krawitz, Kim, Busemeyer, & Brown, 2011a; Behrens, Woolrich, Walton, & Rushworth, 2007; Daw et al., 2006; Gläscher et al., 2009; Gläscher, Daw, Dayan, & Doherty, 2010; Hampton et al., 2006; Iglesias et al., 2013; Kable & Glimcher, 2007; O'Doherty et al., 2007; O'Doherty, Critchley, Deichmann, & Dolan, 2003; Xiang et al., 2013) to identify brain regions that presumably implement a cognitive process of interest.

The hBayesDM package allows users to extract various model-based regressors that can be used for model-based fMRI or EEG analysis (see Figure 7). All model-based regressors are contained in the `modelRegressor` element. Note that in the current version (version 0.2.3.2), only the orthogonalized GNG task provides model-based regressors. The hBayesDM package provides the following model-based regressors, and users can convolve these trial-by-trial data with a hemodynamic response function with their favorite package (e.g., in Statistical Parametric Mapping (SPM; http://www.fil.ion.ucl.ac.uk/spm/), users can use the ‘parametric modulation’ command with a model-based regressor):

1. Stimulus value: *V_t_* (*s_t_*) (stored as `SV`; available in `gng_m3` and `gng_m4`)
2. Action value: *Q_t_* (*go*) (stored as `Qgo`) and *Q_t_* (*NoGo*) (stored as `Qnogo`)
3. Action weight: *W_t_* (*go*) (stored as `Wgo`) and *W_t_* (*NoGo*) (stored as `Wnogo`)

**Figure 7.**
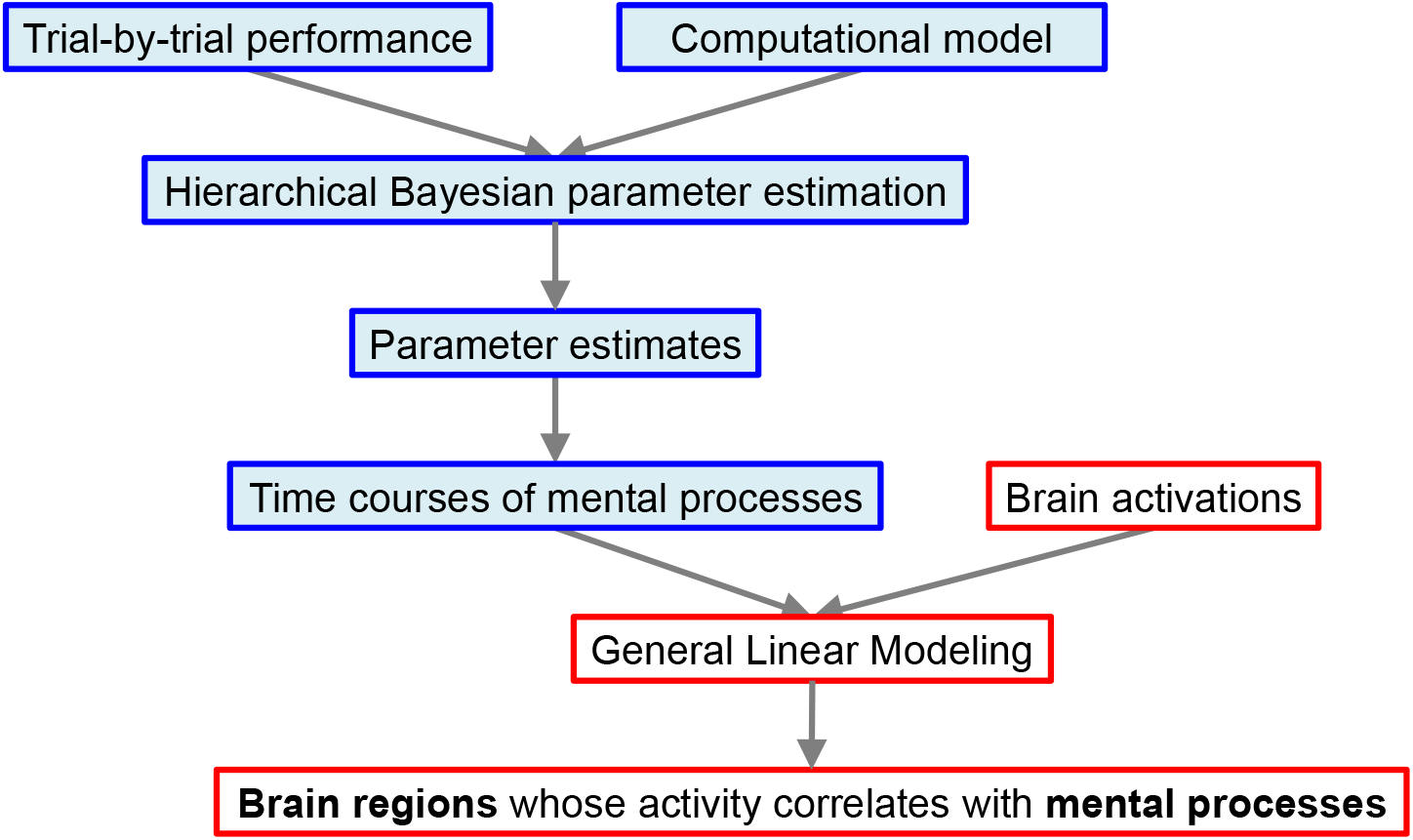
Steps of model-based fMRI. With the hBayesDM package, users can perform the steps highlighted in blue. Users need to use a neuroimaging tool of their choice (e.g., SPM) to perform steps highlighted in red.

For example, to retrieve the stimulus value (=*V_t_* (*s_t_*)) of the group 1 in the previous example (output is saved as `output_group1`), type:

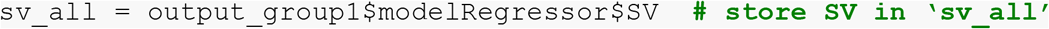

Here, `sv_all` is an array (the number of rows = the number of subjects & the number of columns = the number of trials). Similarly, to retrieve action weight values (*W_t_* (*go*) and *W_t_* (*NoGo*)), type:

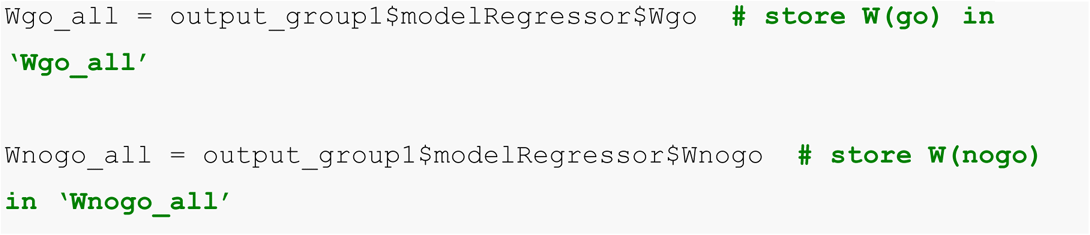

Users can use these values for each subject to perform model-based fMRI analysis with their favorite neuroimaging package (O'Doherty et al., 2007). Once model-based regressors are entered as parametric modulators in the GLM, neuroimaging tools convolve the regressors with the HRF and construct a GLM. For step-by-step tutorials for model-based fMRI, see the following online documents (http://www.translationalneuromodeling.org/uploads/Mathys2016_SPMZurich_ModelBasedfMRI.pdf; http://www.translationalneuromodeling.org/uploads/DiaconescuAndreea_Model-based_fMRI.pdf; http://www.srndna.org/conference2015/files/2014/11/SRNDNA_RL_Modeling_wkshp2.pdf).

## 6. Future directions

In the current version, the hBayesDM package selectively implements eight commonly used RLDM tasks and their models, but we plan to expand the list of tasks and models, so that the hBayesDM can handle an extensive list of RLDM tasks. Latent model-based regressors are available only for a single task, but they will be available for more tasks in the hBayesDM package in a future release. We also plan to develop a GUI interface using a Shiny framework (https://shiny.rstudio.com/) so that users can select a dataset and run models without any R programming.

The hBayesDM package is useful for researchers across all level of experience including experts in computational modeling – hBayesDM systematically implements HBA of various computational models and we find it useful and easier to build new models based on the existing framework. We welcome collaboration and others’ contributions to the package. We plan to release a more detailed tutorial on how to modify existing codes and build new models based on our framework.

In our HBA framework, it is assumed that there is a single hyper-group across all subjects. While it allows more precise estimates with a modest number of subjects (Ahn, Krawitz, Kim, Busemeyer, & Brown, 2011a; Katahira, 2016), the assumption might be invalid with a large (e.g., ˜1000) number of subjects (Ahn & Busemeyer, 2016; Ratcliff & Childers, 2015). Bayesian hierarchical mixture approaches (Bartlema, Lee, Wetzels, & Vanpaemel, 2014) or HBA on subgroups first clustered by behavioral indices (Ahn et al., in preparation) might be an alternative solution when we need to fit large number samples.

In conclusion, the hBayesDM package will allow researchers with minimal quantitative background to do cutting-edge hierarchical modeling on a variety of RLDM tasks. With hBayesDM, researchers can also easily generate model-based regressors required for model-based fMRI/EEG analysis. It is our expectation that the hBayesDM package will contribute to the dissemination of computational modeling and computational psychiatric research to researchers in various fields including mental health.

## Acknowledgements

W.-Y.A. programmed prototypes for several tasks while he was advised by Jerome Busemeyer (for the Iowa Gambling Task) or P. Read Montague/Peter Dayan (for Orthogonalized Go/NoGo, Two-Step, and Risk-Aversion tasks as well as the Ultimatum Game). We thank them for their guidance and resources provided to W.-Y.A. We thank Peter Sokol-Hessner for sharing data published in Sokol-Hessner et al. (2009). L.Z. was partially supported by German Research Foundation (DFG GRK 1247) and Bernstein Computational Neuroscience Program of the German Federal Ministry of Education and Research (BMBF Grant 01GQ1006).

### Author contributions

W.-Y.A. conceived and designed the project. W.-Y.A., N.H., and L.Z. programmed codes for hierarchical Bayesian modeling. N.H. built an R package and wrote help files. W.-Y.A., N.H., and L.Z. wrote the paper.

